# Epigenetic changes associated with hyperglycaemia exposure in the longitudinal D.E.S.I.R. cohort

**DOI:** 10.1101/2022.03.18.484909

**Authors:** Amna Khamis, Lijiao Ning, Beverley Balkau, Amélie Bonnefond, Mickaёl Canouil, Ronan Roussel, Philippe Froguel

## Abstract

**Aim:** Understanding DNA methylation dynamics associated with progressive hyperglycaemia exposure could provide early diagnostic biomarkers and an avenue for delaying type 2 diabetes (T2D) disease. We aimed to identify DNA methylation changes during a 6-year period associated with early hyperglycaemia exposure using the longitudinal D.E.S.I.R. cohort.

**Methods:** We selected individuals with progressive hyperglycaemia exposure based on T2D diagnostic criteria: 27 with long-term exposure, 34 with short-term exposure and 34 normoglycaemic controls. DNA from blood at inclusion and at the 6-years visit was subjected to methylation analysis using 850K methylation-EPIC arrays. A linear mixed model was used to perform an epigenome-wide association study (EWAS) and identify methylated changes associated with hyperglycaemia exposure during 6-year time-period.

**Results:** We did not identify differentially methylated sites that reached FDR-significance in our cohort. Based on EWAS, we focused our analysis on methylation sites that had a constant effect during the 6-years across the hyperglycaemia groups compared to controls and found the most statistically significant site was the reported cg19693031 probe (*TXNIP*). We also performed an EWAS with HbA1c, using the inclusion and the 6-years methylation data and did not identify any FDR-significant CpGs.

**Conclusions:** Our study reveals that DNA methylation changes are not robustly associated with hyperglycaemia exposure or HbA1c during a short-term period, however, our top *loci* indicate potential interest and should be replicated in larger cohorts.

## 1. Introduction

Type 2 diabetes (T2D) is a progressive disease that is often asymptomatic for many years prior to diagnosis [1], thereby, disease management can be improved through the identification of early biomarkers. Human and animal studies have demonstrated that epigenetics may play a role in maintaining a metabolic memory in response to the environmental exposure of modest hyperglycaemia [2]. For instance, *in vitro* studies in animal models have demonstrated that poor glucose control during early-life was associated with long-term epigenetic changes that persisted over the lifespan, even following glucose lowering treatment [3], [4]. In human studies, we have shown that methylation sites in blood DNA in genes were associated with incident T2D in the multi-ethnic LOLIPOP cohort [5], *i.e*., epigenetic markers could predict incident T2D. Since then, other studies have identified methylated DNA sequences (CpGs) associated with T2D in nearby candidate genes with defined biological function, including glycolysis (*SREBF1*), glucose homeostasis (*TXNIP*) and lipid metabolism and insulin resistance (*ABCG1*) [5]–[7]. Collectively, these studies demonstrate that identifying early mechanisms associated with T2D risk may provide opportunities for early interventions and lead to better management of T2D. However, most of these studies were cross-sectional, in individuals following T2D diagnosis or using one time-point, therefore, the behaviour of these epigenetic changes over time associated with early hyperglycaemic exposure has not been explored. Our aim was to quantify this DNA-based metabolic memory by studying the relationship between methylation and exposure to hyperglycaemia in response to long-term (> 6 years) and short-term (3-6 years) hyperglycaemia, in addition to HbA1c, using the longitudinal French D.E.S.I.R. population-based cohort (study design is summarised in Figure 1).

**Figure 1:**
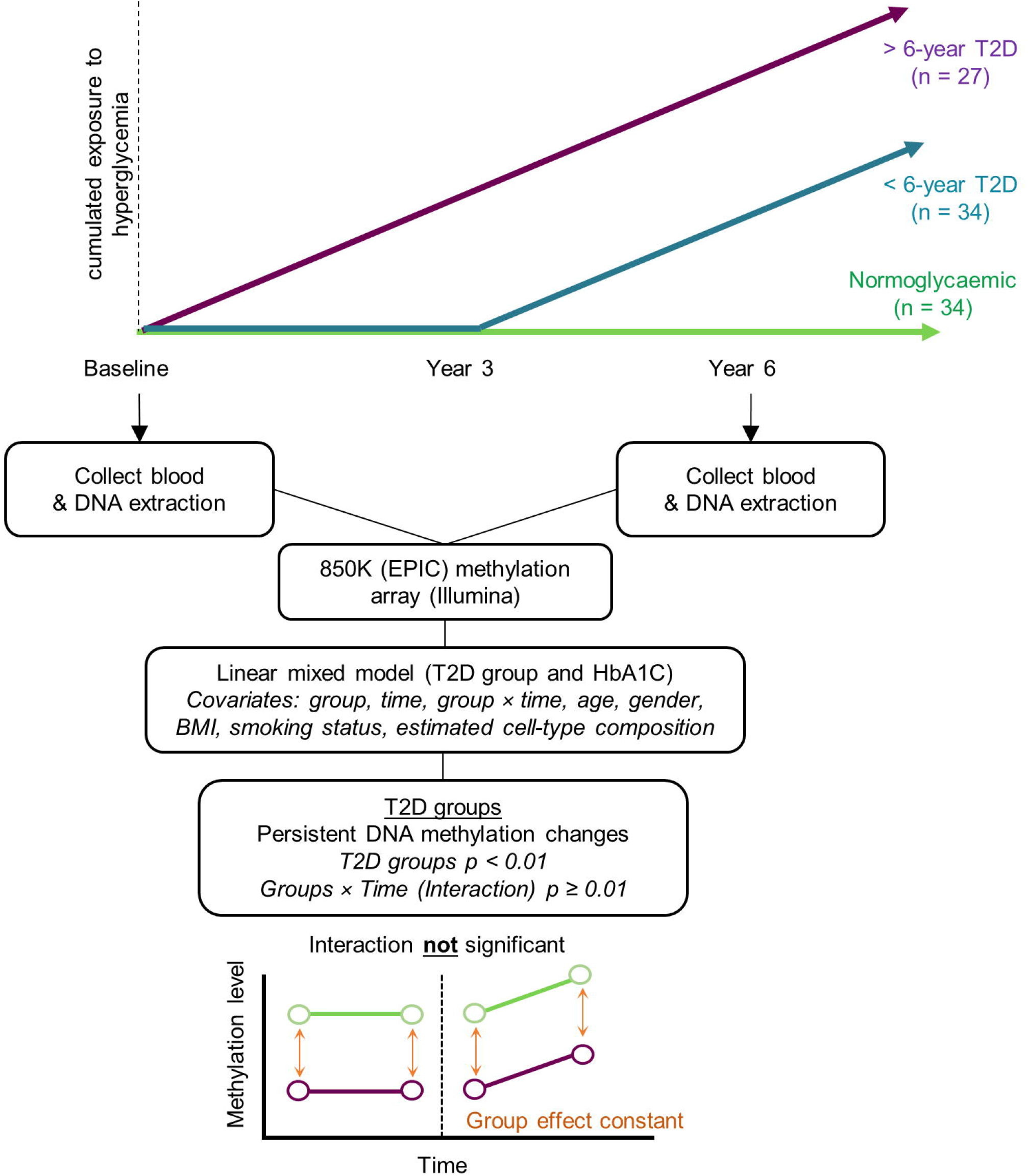
Summary of the study design. Individuals were selected based on three cumulated exposure to hyperglycaemia: those with long-term hyperglycaemia, screened with T2D at baseline (with exposure to hyperglycaemia and increasing cumulated exposure during the 6-year period), those with short-term hyperglycaemia, screened with T2D at 3 years from baseline (which starts at some point between baseline and 3 years, and then accumulates until 6 years) and normoglycaemic controls. DNA collected from whole blood at baseline and at the 6-year follow-up was subjected to whole methylome analysis. An EWAS was performed to identify differentially methylated sites associated with hyperglycaemia exposure. A focused analysis was performed to identify differentially methylated sites with a statistically significant difference between exposed and unexposed groups, constant at the two time-points. For simplicity, this illustration demonstrates only one exposure group.

## 2. Materials and methods

### Cohort

We used the D.E.S.I.R. (Data from an Epidemiological Study on the Insulin Resistance Syndrome) cohort [8], [9], which is a French-based cohort including adults aged between 30 and 50 years recruited from 10 recruitment centres in France. Participants were followed up every 3 years for 9 years, where detailed clinical measurements, including fasting glucose and glycated haemoglobin A1c (HbA1c), were collected at all time-points. All participants signed an informed consent, and the protocol was approved by the ethics committee of Kremlin Bicêtre Hospital in Paris, France. All experiments and methods presented in this study were performed in accordance with relevant guidelines and regulations.

We selected individuals in our study with available whole blood samples collected at baseline and at the 6-year follow-up and that matched in age, sex and body mass index (BMI) in three progressive glycaemic exposure groups. These groups were defined based on the American Diabetes Association (ADA) guidelines [10], which uses an HbA1c ≥ 6.5 % or a fasting plasma glucose ≥ 7.0 mmol/L. This resulted in the following groups: 1) the long-term exposure group included 27 participants with T2D from baseline and had a cumulative hyperglycaemia exposure for > 6 years at the 6 year visit, 2) the short-term exposure group included 34 participants with T2D by the 3-year follow-up, *i.e*., not yet diagnosed at baseline and therefore had 3-6 years hyperglycaemia exposure by the 6-year follow-up, and 3) 34 normoglycaemic participants who were non-diabetic at all follow-up visits. For simplicity, hyperglycaemia groups will be referred to as short-term and long-term exposure. The trajectory of the fasting glucose and HbA1c levels are shown in Supplementary Figure A. A summary of the cohort characteristics is shown in Table 1.

**Table 1:** Clinical characteristic of cohort; The D.E.S.I.R. Study

|  | Normoglycaemic Control<sup>1</sup><br>(N = 33) | Short-term Exposure<sup>1</sup><br>(N = 34) | Long-term Exposure<sup>1</sup><br>(N = 27) | P-value<sup>2</sup> |
| --- | --- | --- | --- | --- |
| Age at baseline (years) | 51.64 ± 8.48 | 50.94 ± 8.14 | 54.44 ± 7.30 | 1.7 × 10<sup>-1</sup> |
| BMI at baseline (m/kg<sup>2</sup>) | 28.02 ± 3.94 | 29.55 ± 3.79 | 27.94 ± 3.24 | 1.5 × 10<sup>-1</sup> |
| Gender |  |  |  | 1 |
| Female | 9 (27%) | 9 (26%) | 8 (30%) |  |
| Male | 24 (73%) | 25 (74%) | 19 (70%) |  |
| Smoking at baseline |  |  |  | 3.0 × 10<sup>-1</sup> |
| Never | 14 (42%) | 11 (32%) | 14 (52%) |  |
| Other | 19 (58%) | 23 (68%) | 13 (48%) |  |
| HbA1c at baseline (%) | 5.41 ± 0.30 | 5.85 ± 0.32 | 6.61 ± 1.17 | 5.2 × 10<sup>-11</sup> |
| HbA1c at 6-year visit (%) | 5.46 ± 0.25 | 6.37 ± 0.90 | 6.84 ± 1.21 | 1.2 × 10<sup>-10</sup> |
| Fasting glucose at baseline (mmol/L) | 5.11 ± 0.30 | 6.20 ± 0.48 | 8.27 ± 1.99 | 1.3 × 10<sup>-1</sup> |
| Fasting glucose at 6-year visit (mmol/L) | 5.08 ± 0.32 | 7.29 ± 1.70 | 7.79 ± 2.18 | 7.8 × 10<sup>-14</sup> |
| Glucose lowering drugs at baseline = No | 0 (0%) | 0 (0%) | 0 (0%) | 1 |
| Glucose lowering drugs at 6-year visit |  |  |  | 9.3 × 10<sup>-6</sup> |
| Yes | 0 (0%) | 11 (34%) | 12 (46%) |  |
| Missing | 0 | 2 | 1 |  |
| <sup>1</sup>Statistics presented: n (%) for discrete variables and mean ± SD for continuous variables.<br><sup>2</sup>Comparison using Fisher's exact test for discrete variables and Kruskal-Wallis Rank Sum test for continuous variables. |  |  |  |  |

### Whole methylome analysis

Whole blood was collected from 95 participants at two time-points, at baseline and at the 6-year visit. DNA was extracted using the DNeasy Blood & Tissue kit (Qiagen, Germany). All DNA samples were subjected to DNA methylation profiling. Briefly, bisulfite conversion of genomic DNA was performed using the EZ-96 DNA Methylation kit (Zymo Research) following the manufacturer’s protocols. Bisulfite-converted DNA was assayed for DNA methylation using Illumina’s Infinium 850K MethylationEPIC array. Data analyses were performed using the R software (version 4.0.3). The DNA methylation IDAT files were imported using the R package *minfi* [11] for pre-processing and quality-control. The following probes were excluded from further analysis: probes with a detection *p-*value ≥ 0.01 in at least one sample, cross-hybridizing probes, probes with a bead count less than 3 in at least 5 % of the samples, non-CpG probes and probes which lie near single nucleotide polymorphisms (SNPs). Of the 866,895 probes in the array, a total of 148,306 were excluded. Probes on sex chromosomes were used for sex estimation and excluded from downstream analyses. A comparison between phenotype and estimated sex based on DNA methylation identified one sample with a sex discrepancy, which was excluded from further analyses. Samples with less than 95 % of probes with a detection *p*-value < 0.01 were excluded. Probe type design biases and batch effects were corrected using the R package *Enmix* [12]. After quality-control, 94 samples and 718,589 CpG probes were used for further downstream analyses. As blood samples are expected to include a variety of cell types and which might have a potential confounding effect on DNA methylation, we estimated cell composition using a whole blood reference panel from the R package *FlowSorted.Blood.EPIC* [13]. Methylation levels are denoted by β-values (where 0 indicates 0 % methylation and 1 indicates 100 % methylation) and transformed into M-values for analyses [14].

### Statistical analyses

To test our hypothesis that epigenetic changes were associated with hyperglycaemic exposure, we performed an EWAS using a linear mixed model performed using R packages *lme4* [15] and *lmerTest*[16], adjusted for age, sex, BMI, smoking status, time-point and estimated cell-type composition. The EWAS was conducted to identify methylation changes associated with glycaemic exposure in our groups (*i.e*., long-term *vs*. control, short-term *vs*. control and long-term *vs*. shortterm). An interaction term between time-point and glycaemic exposure group was included in the model. Sample-level intercept was included to take into account within sample correlation across measurements. P-values were corrected for multiple testing using the false discovery rate (FDR) method from Benjamini-Hochberg. An ANOVA was applied on the mixed model to evaluate the overall effect of group and the p-value of ANOVA test was used to rank CpGs. Based on EWAS, we focused on the sets of differentially methylated sites which epigenetic changes in hyperglycaemia exposure groups were statistically different from the normoglycaemic controls and this effect is constant across two time-points, where the group effect p-value < 0.01 and an interaction with p-value ≥ 0.01 were considered.

For association with HbA1c, an EWAS using a linear mixed model was performed, taking into account the HbA1C values from baseline and year 6 for each individual. The model included age, sex, BMI, smoking status, time-point and estimated celltype composition as covariates. P-values were corrected for multiple testing by Benjamini-Hochberg’ FDR method. Prior to the analysis, we identified four individuals with HbA1c values considered as potential outliers (1.5 times the interquartile range of the average HbA1c of two time-points), which were removed from further analysis.

## 3. Results

We performed an EWAS on individuals with short-term and long-term exposure, compared to controls to identify differentially methylated sites associated with progressive hyperglycaemic exposure. We did not observe any FDR-significant methylated sites associated with hyperglycaemia exposure for short-term *vs*. control, long-term *vs*. control or long-term *vs*. short-term hyperglycaemia exposure (Figure 2).

**Figure 2:**
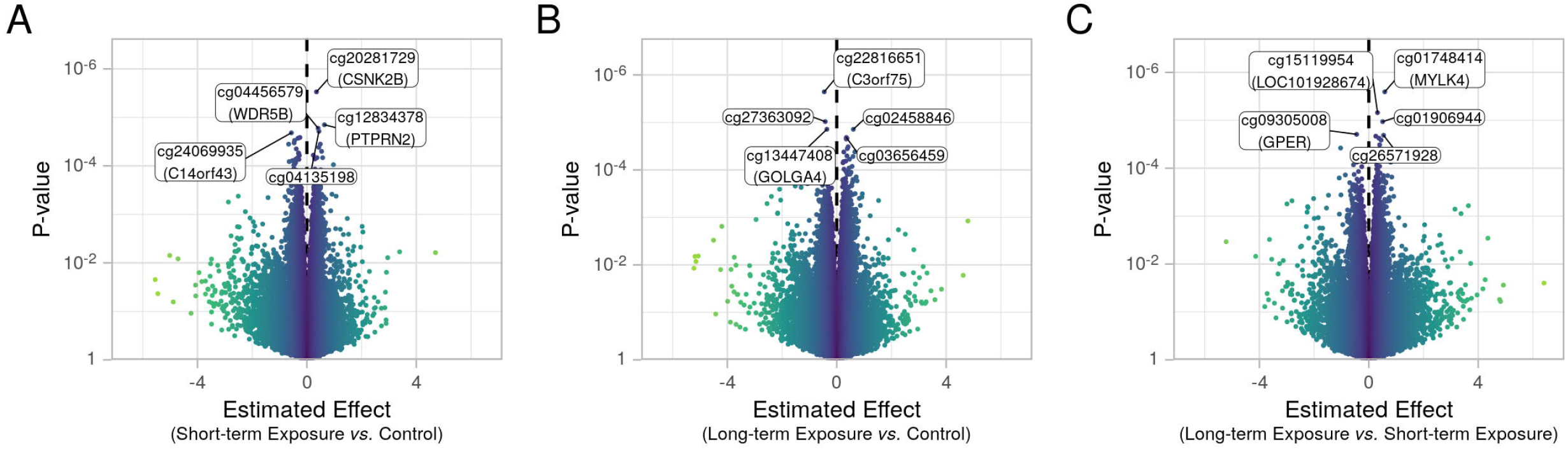
EWAS with hyperglycaemic exposure: Volcano plots for the three comparisons and highlighting the 5 most significant CpGs.

We then focused our analysis to identify methylation changes that had a longitudinal constant effect in the two hyperglycaemic groups compared to controls. We found that the most differentially methylated CpG was at the cg19693031 probe, located at the 3’UTR of the *Thioredoxin Interacting Protein* (*TXNIP*) gene (Figure 3a). The probe was hypomethylated in both hyperglycaemic exposed groups compared to the normoglycaemic group (short-term exposure group: p-value = 2.6 × 10^−4^; long-term exposure group: p-value = 9.1 × 10^−5^; no significant difference between the two exposed groups: p-value = 6.6 × 10^−1^). The five most statistically significant associations are shown in Figure 3. In total, we found a total of 216 differentially methylated sites that had a constant effect at baseline and at the 6-year follow-up, distributed in 145 unique genes and 64 in intergenic regions (Appendix A). To further elucidate the biological relevance of the identified DMPs, we performed a literature search to identify a potential role in T2D and glycaemia. We found that 16 % of the 145 differentially methylated sites annotated to genes with an already reported role in diabetes, glycaemia or islet function (Appendix B). This included genes with single nucleotide polymorphisms (SNPs) reported to be associated with T2D, such as *PLA2G6* [17] (Figure 4a), in addition to others with a role in insulin secretion in pancreatic islets, such as *SORCS1* [18] (Figure 4b). We found a site within the 5’UTR of the *FAM123C* gene, where epigenetic changes in the 5’UTR of the same gene have been previously reported to be associated with age in both islets and whole blood [19] (Figure 4c).

**Figure 3:**
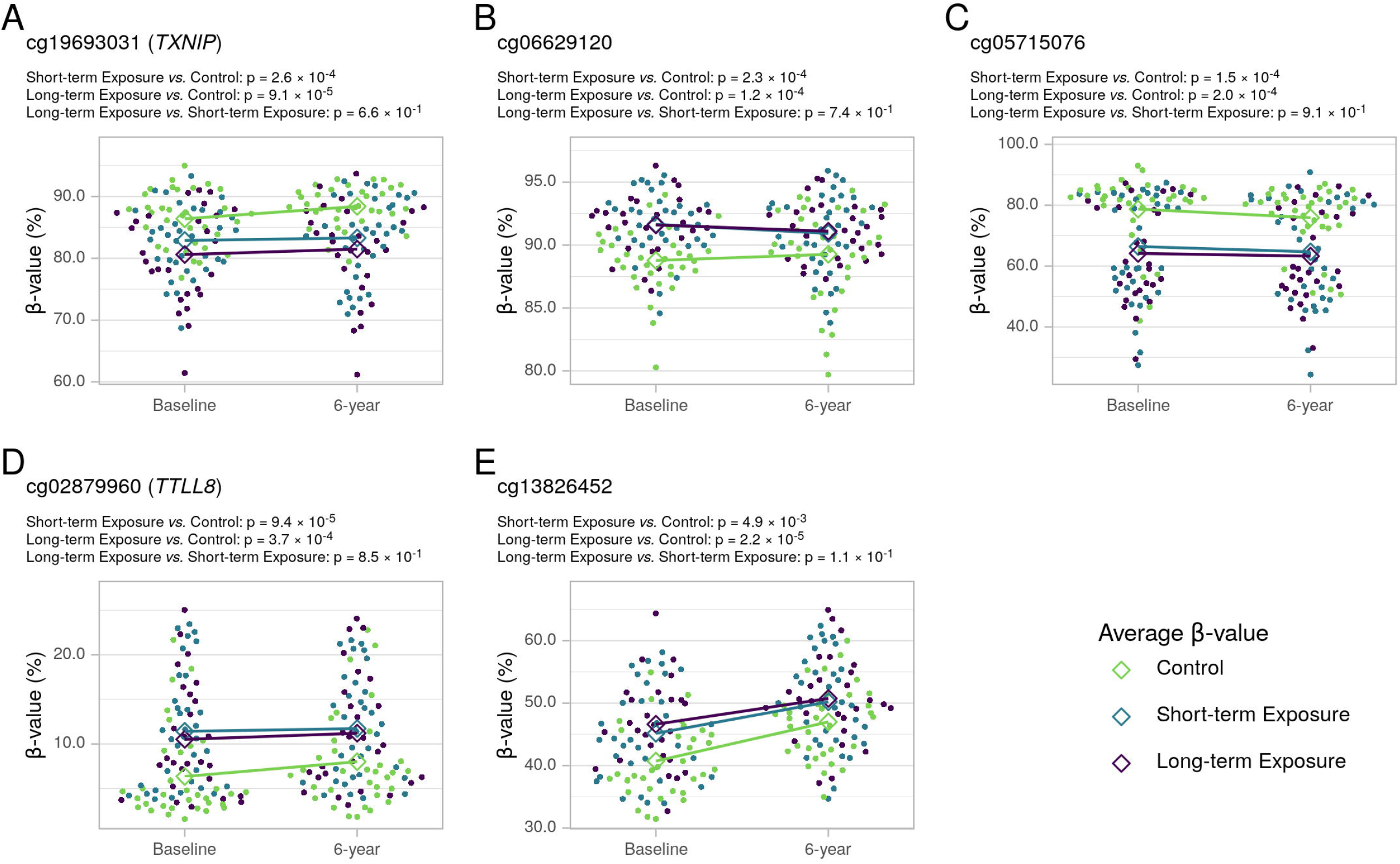
Most significant methylation changes persistent at baseline and 6-year follow-up. Scatter plot of methylation levels of the five CpGs with the most statistically significant differences between the exposed groups and the normoglycaemic control group. P-values are shown comparing each hyperglycaemia exposure group to the control group and between the two exposure groups. The lines connect the intragroup average β-value at the two time-points.

**Figure 4:**
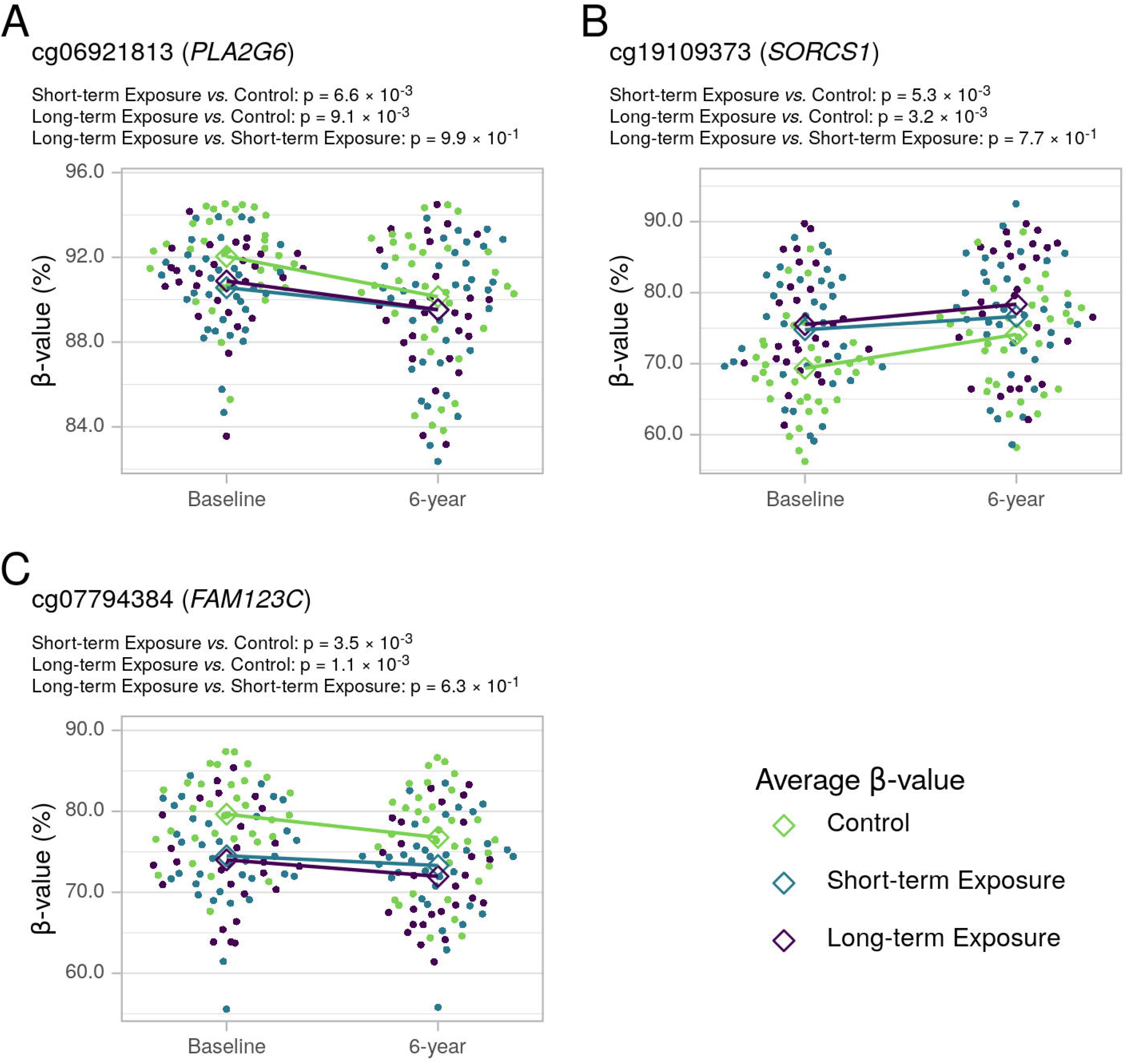
Methylation changes in genes with a known role in T2D, glycaemic and islet function. Scatter plot of methylation levels of genes associated with a) T2D GWAS, such as *PLA2G6* and b) genes involved in insulin secretion, such as *SORCS1* and c) methylation changes in *FAM123C* that have been found to be associated with ageing in pancreatic islets. P-values for the exposure groups compared to the control group and between two exposure groups are presented. The lines connect the intragroup average β-value at the two time-points.

We performed an EWAS using a linear mixed model to identify methylation changes associated with HbA1c as a quantitative trait. We did not identify any FDR-significant associations with HbA1c. The most significant differentially methylated CpGs were located in the body of the *SECISBP2* gene (p = 1.65 × 10^−6^), the TSS1500 of the LOC100128531 (p = 4.53 × 10^−6^), and the body of the antisense gene MYLK-AS1 (p = 5.06 × 10^−6^) (Figure 5; Appendix C).

**Figure 5:**
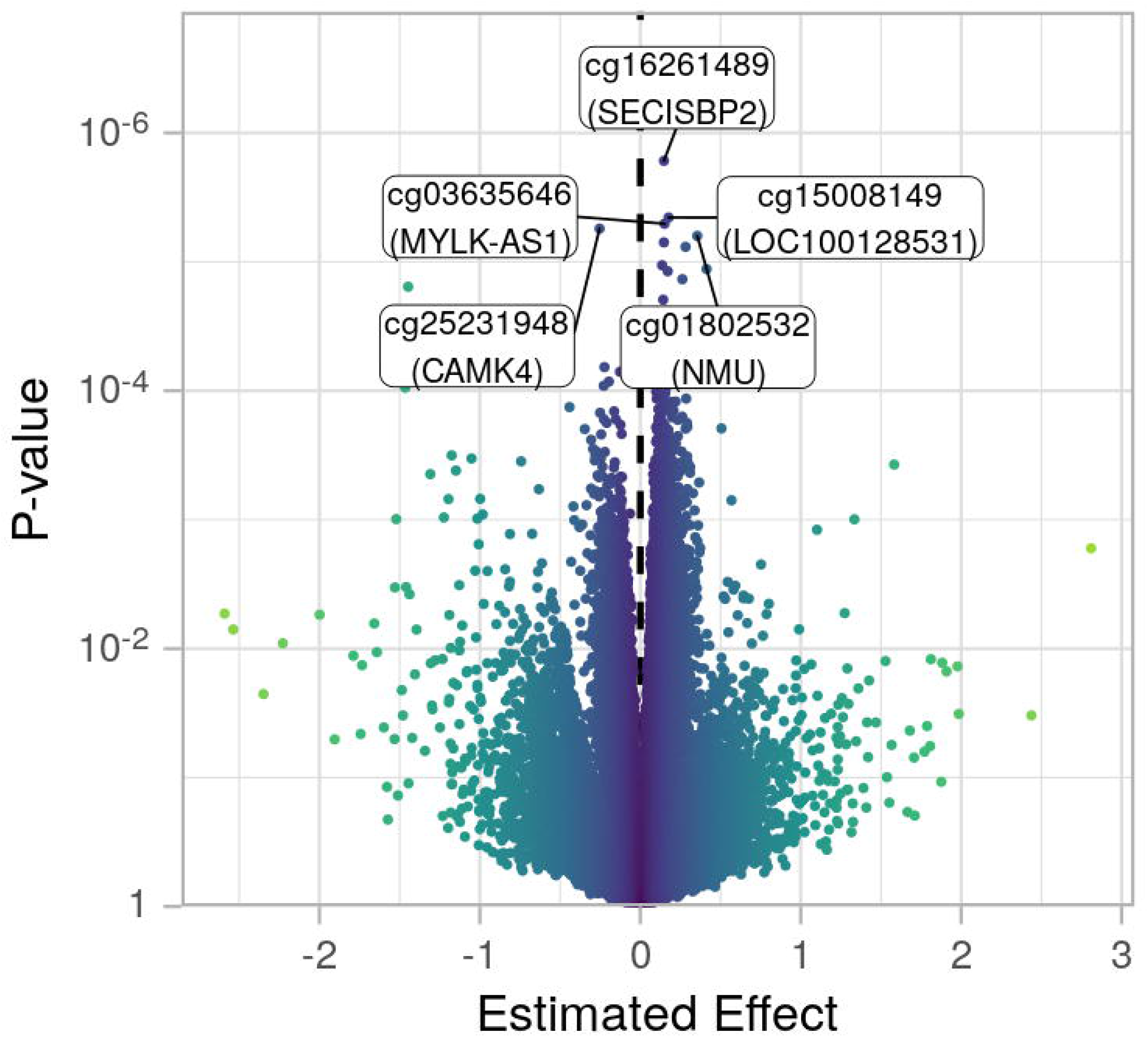
Most significant methylation changes associated with HbA1c. A volcano plot for the EWAS results and highlighting the 5 most significant CpG.

## 4. Discussion

Here, we explored the blood methylation dynamics during a 6-year period associated with hyperglycaemia exposure in a total of 96 individuals using the D.E.S.I.R. longitudinal cohort. Although cross-sectional studies have identified several CpG sites associated with T2D and HbA1c [5], [20], [21], we did not identify any differentially methylated sites following multiple testing correction using both T2D progression groups and HbA1c level during a 6-year period, demonstrating that in our study, methylation changes do not show noticeable glucose-dependent changes during the early phases of hyperglycaemic exposure. However, our focused analysis demonstrates that the majority of CpGs showed similar levels between the two hyperglycaemia exposure groups at baseline and 6-year follow-up, suggestive of a history of differential methylation processes not limited to early response to hyperglycemia.

There are several indications to support the robustness of our method. For instance, differential methylation in several of our top most significant *loci* have already been established within the context of T2D. In our focused analysis, the methylation site that differed the most from the normoglycaemic group was the cg19693031 probe located at the *TXNIP* gene, confirming several studies that found this probe to be associated with T2D, HbA1c and associated traits [6], [22]–[25]. Moreover, an almost two-decade longitudinal study found that changes in *TXNIP* CpGs in type 1 diabetes were associated with an increased risk of later development of diabetic complications [26], [27]. In addition, in a recent study, we found a hypermethylation of two further probes in the *TXNIP* gene that were associated with fetal exposure of area under the curve of glucose during pregnancy [28], which was associated with T2D later in life and correlated with *TXNIP* gene expression in liver. Indeed, these studies demonstrate that this CpGs is differentially methylated prior to the diagnosis of established T2D (fasting glucose > 7 mmol/L), highlighting its potential importance as a predictor of T2D future development in early prediabetes states.

Additionally, we found that one of our most significant *loci* associated with T2D groups was located in the *PTPRN2* gene. A recent study identified a hypermethylation in this gene was differentially methylated in pancreatic islets of a pre-diabetes mouse model [28]. The authors further confirmed a *PTPRN2* hypermethylation was one of the strongest blood-based T2D predictive marker in the human (EPIC)-Potsdam cohort (median 3.8 years prior to T2D diagnosis) and also hypermethylated in pancreatic islets from T2D individuals [28]. Indeed, our study found that a hypermethylation, also located in the gene body of *PTPRN2* gene, was the second most significant differentially methylated site in our 3-6 years *vs*. control EWAS (Figure 2a). Therefore, this suggests that although our study was performed in whole blood, a tissue not directly related to T2D, these mechanisms potentially have a role in target pancreatic islet tissues, indicating that these *loci* may mirror changes in crucial biologically relevant tissue.

We present a unique study and statistical method for identifying differentially methylated *loci* in a longitudinal study design. Indeed, this design and the central focus on blood-based DNA methylation signatures in response to hyperglycaemic exposure allowed us to explore differentially methylated sites that would have been missed in already published cross-sectional studies. Moreover, it is essential to note that this study has several limitations that are important to consider. First, we were only able to evaluate two time-points and are, therefore, unable to conclusively and precisely determine the dynamics of these methylation sites within the 6-year duration. In that context, the 6-year duration is a short-term period and it is possible that a longer duration is necessary to capture these differences. In addition, the number of participants in our study sample is modest, and larger studies are still needed to confirm the associations we have reported here and to further investigate their biological function.

## 5. Conclusion

In conclusion, we did not find any robust CpGs associated with glycaemic measures within a 6-year period of T2D progression. However, our nominal results identified several *loci* that are consistent between T2D groups (short-term and long-term) and controls, which could potentially have a predictive value and should be further replicated in larger cohorts with longer duration between measurements to fully capture these differences.

## Abbreviations

D.E.S.I.R.: Data from an Epidemiological Study on the Insulin Resistance Syndrome
EWAS: Epigenome-wide association study
FDR: false discovery rate
HbA1C: glycated haemoglobin A1c
SNP: single nucleotide polymorphism
T2D: type 2 diabetes

## Acknowledgements

The authors would like to thank Stephane Lobbens, Nicolas Larcher and Frédéric Allegaert for their technical support. The D.E.S.I.R. study has been funded by INSERM contracts with Caisse Nationale de l’Assurance Maladie des Travailleurs Salariés (CNAMTS), Lilly, Novartis Pharma, and Sanofi-Aventis; INSERM (Réseaux en Santé Publique, Interactions entre les detérminants de la santé, Cohortes Santé TGIR 2008); the Association Diabète Risque Vasculaire; the Fédération Française de Cardiologie; La Fondation de France; Association de Langue Française pour l’Etude du Diabète et des Maladies Métaboliques (ALFEDIAM)/Société Francophone de Diabétologie (SFD); l’Office National Interprofessionnel des Vins (ONIVINS); Ardix Medical; Bayer Diagnostics; Becton Dickinson; Cardionics; Merck Santé; Novo Nordisk; Pierre Fabre; Roche; Topcon.

## The D.E.S.I.R. Study Group

CESP, Inserm U1018: B. Balkau, P. Ducimetière, E. Eschwège; Univ Paris Descartes: F. Rancière; Inserm U367: F. Alhenc-Gelas; CHU d’Angers: A. Girault; Bichat Hospital: F. Fumeron, M. Marre, R Roussel; CHU de Rennes: F. Bonnet; CNRS UMR8090, Lille: A. Bonnefond, S. Cauchi, P. Froguel; Centres d’examens de santé de l’Assurance Maladie: Alençon, Angers, Blois, Caen, Chateauroux, Chartres, Cholet, Le Mans, Orléans, Tours; Institut de Recherche en Médecine Générale: J. Cogneau; General practitioners of the Region; Institut inter-Régional pour la Santé (IRSA): C. Born, E. Caces, M. Cailleau, O Lantieri, J.G. Moreau, F. Rakotozafy, J. Tichet, S. Vol.

## Availability of data and materials

The data underlying the results presented in the study are available upon publication.

## Authors’ contributions

P.F. and R.R. designed the study. M.C., L.N. and A.K. have performed the statistical analyses and generated the associated figures and tables. A.B, R.R. and P.F. contributed to data. A.K. has interpreted data and has written the manuscript. B.B. designed and managed the D.E.S.I.R. study. All authors critically reviewed and edited the manuscript.

