## Supplementary figures and images for "Epigenetic changes associated with hyperglycaemia exposure in the longitudinal D.E.S.I.R. cohort"

### Supplementary Figure A

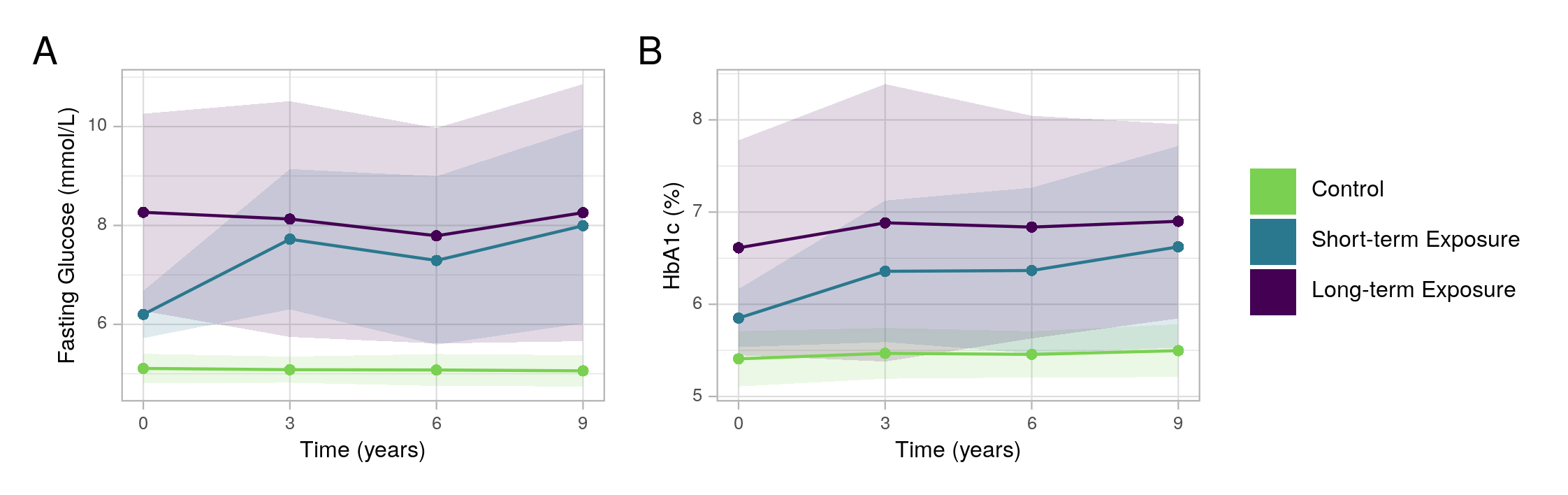
